# Programmable delivery of fluoxetine via wearable bioelectronics for wound healing in vivo

**DOI:** 10.1101/2023.10.10.561754

**Authors:** Houpu Li, Hsin-ya Yang, Narges Asefifeyzabadi, Prabhat Baniya, Andrea Medina Lopez, Anthony Gallegos, Kan Zhu, Hao-Chieh Hsieh, Tiffany Nguyen, Cristian Hernandez, Ksenia Zlobina, Cynthia Recendez, Maryam Tebyani, Héctor Carrión, John Selberg, Le Luo, Moyasar A. Alhamo, Athena M. Soulika, Michael Levin, Narges Norouzi, Marcella Gomez, Min Zhao, Mircea Teodorescu, Roslyn Rivkah Isseroff, Marco Rolandi

## Abstract

The ability to deliver drugs with precise dosages at specific time points can significantly improve disease treatment while reducing side effects. Drug encapsulation for gradual delivery has opened up the doors for superior treatment regimen. To expand on this ability, programming bioelectronic devices to deliver small molecules enables ad-hoc personalized therapeutic profiles that are more complex than simple gradual release. Here, we introduce a wearable bioelectronic bandage with an integrated electrophoretic ion pump that affords on-demand drug delivery with precise dose control. Delivery of fluoxetine to wounds in mice resulted in a 27.2% decrease in the macrophage ratio (M1/M2) and a 39.9% increase in re-epithelialization, indicating a shorter inflammatory phase and faster overall healing. Programmable drug delivery using wearable bioelectronics in wounds introduces a broadly applicable strategy for the long-term delivery of a prescribed treatment regimen with minimal external intervention.

## 1 Introduction

The ability to tailor the therapeutic levels of drugs over time to the dynamic evolution of our physiology improves treatment effectiveness and leads to better clinical outcomes [1]. Research in drug delivery has developed materials and chemical strategies for passive and active controlled release [2]. Merging this active control strategy with programmable electronic devices can open the doors to personalized and more effective treatment regimes [3]. Bioelectronic devices are capable of delivering on-demand physiological-active ions [4], small molecules [3b, 5], and electric fields as treatments [6]. This ability is particularly important when applied to rapidly changing environments, such as the one found in wounds [7]. Wound healing usually involves four overlapping phases: hemostasis, inflammation, proliferation, and remodeling [8]. A delay or disruption in one of these phases leads to chronic wounds, scarring, infection, sepsis, and even death [9]. To facilitate wound healing, wearable bioelectronic bandages have employed external stimuli, such as electric fields [10] and biochemicals [11]. To expand on these treatments, here we have developed a programmable bioelectronic bandage capable of delivering the drug fluoxetine (brand name: Prozac) as a personalized treatment regimen to accelerate wound healing in mice (**Figure 1**). Prior studies have demonstrated that topical administration of fluoxetine improves diabetic and non-diabetic wound contraction and closure [12], decreases wound inflammation, and minimizes infection [13].

**Figure 1.**
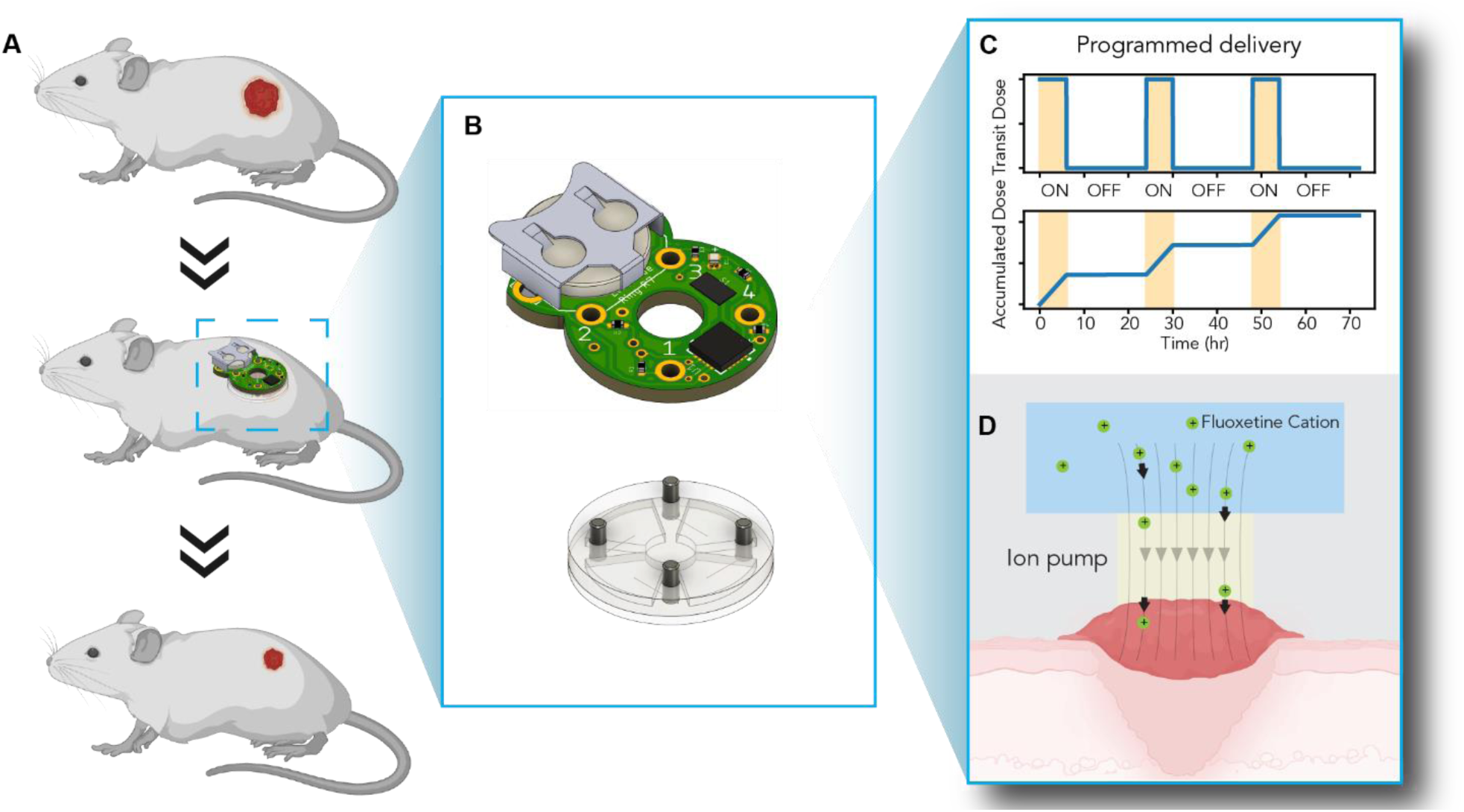
Depicts the wearable bioelectronic bandage designed for programmable drug delivery. (A) A schematic of the wearable bioelectronic bandage applied to a mouse model. (B) A CAD design of the wearable bioelectronic bandage, consisting of a battery-powered PCB controller module (top) and an ion pump made from PDMS (bottom). (C) The transit and accumulated dose curve of the programmed delivery of fluoxetine is plotted. The wearable bioelectronic bandage delivers 30 μg of fluoxetine over 6 hours and repeats daily. (D) Schematics of the ion pumping process show that the electric field drives the fluoxetine cations (green spheres) to the wound through the ion-selective hydrogel (light yellow)

## 2 Results

The bioelectronic bandage is a wearable device (Figure 1) that consists of two modules: an ion pump drug delivery module and a battery-powered controller module (Figure 1B). The controller module is responsible for translating a pre-programmed delivery profile into a sequence of voltage signals (Figure 1C), which activate the ion pump to deliver fluoxetine to the wound (Figure 1D).

The ion pump drug delivery module includes two reservoirs filled with fluoxetine solution, which are connected to the wound bed using two glass capillaries filled with an ion-selective hydrogel (**Figure 2**A). The module is made of polydimethylsiloxane (PDMS) using a casting process (see Materials and Methods, **Figure S1**). Two Ag/AgCl electrodes (a working electrode on the left and a counter electrode on the right) connect the reservoirs to the voltage source on the control module that drives the delivery of fluoxetine using V_Flx_. To allow the delivery of fluoxetine (Flx), the solution in the reservoirs is acidic (0.01M Flx·HCl, tuned to pH 6), making fluoxetine positively charged due to protonation (**Figure S2**). When V_Flx_ is positive, Flx molecules are pushed from the reservoir on the left of the image into the wound bed via the ion-selective hydrogel acting as an ion exchange membrane (IEM) while blocking negatively charged anions entering the reservoirs from the wound bed. To maintain charge neutrality, we presume that physiological cations exit the wound bed through the ion-selective hydrogel on the right and enter the other reservoir that houses the counter electrode. These physiological cations also act as the charge carriers in the wound bed, completing the circuit. The difference in concentration between the physiological cations (∼150 mM) [14] and fluoxetine (<0.3 mM) in the wound bed ensures that most, if not all, of the fluoxetine delivered to the wound bed, does not escape into the reservoir containing the counter electrode.

**Figure 2.**
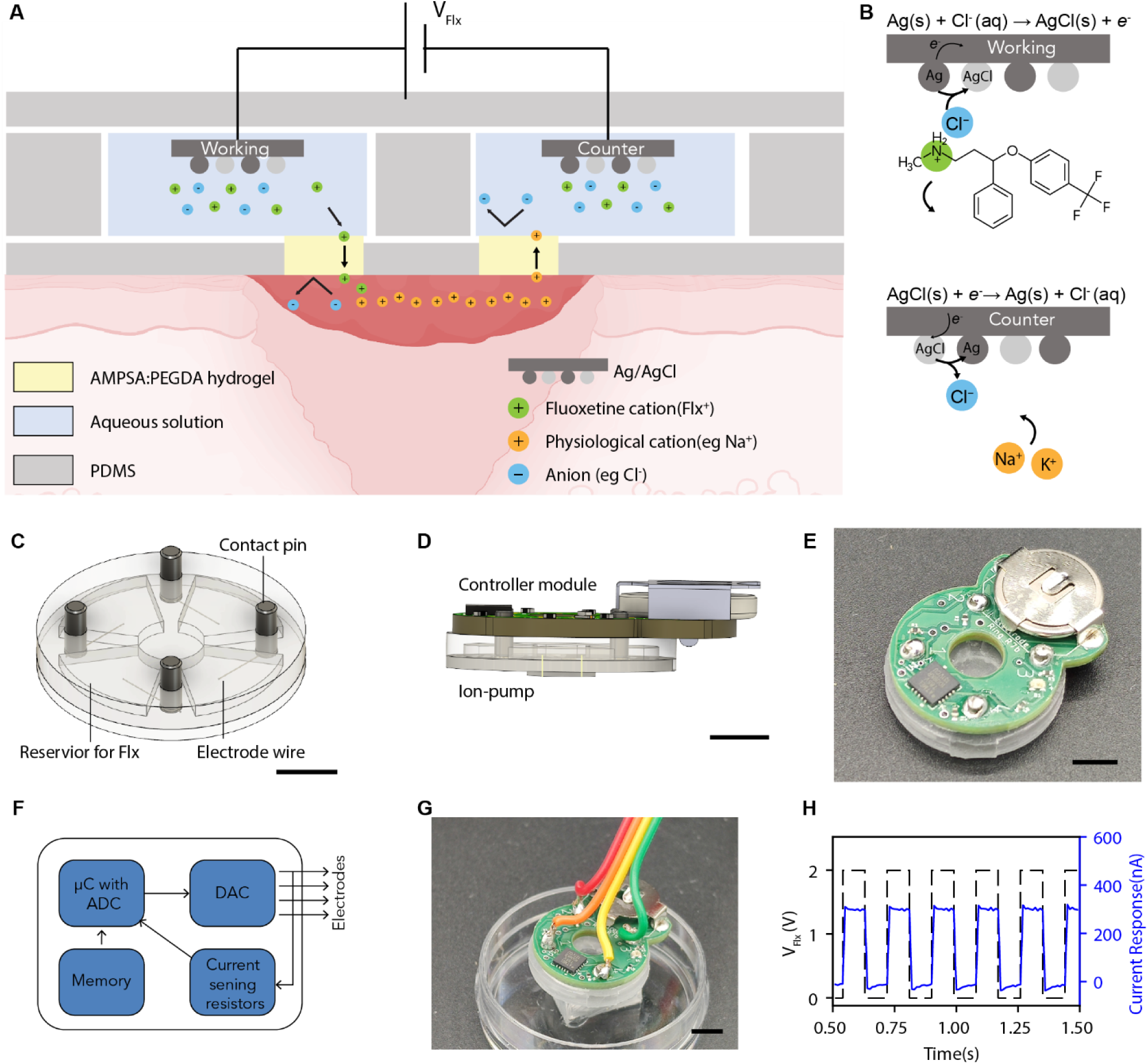
Showcases the design and characterization of the wearable bioelectronic bandages. (A) The ion pump’s working principle is illustrated, where the reservoirs are filled with 0.01 M fluoxetine hydrochloride (FlxCl) solution. Under positive V_Flx_, the Flx^+^ cations migrate from the reservoir to the wound under electrophoretic force and exchange with physiological cations. An ion selective AMPSA:PEGDA polyanion hydrogel (light yellow) is negatively charged, acting as a filter that blocks anions such as Cl^-^, while allowing cations such as Flx^+^ to pass. (B) The electrochemical reaction for the ion pump electrodes is depicted happening at the opposite direction simultaneously. The Ag on the working electrode (gray) becomes oxidized under the electrochemical reaction, absorbs a Cl^-^ from the Flx·HCl solution, releases an electron, and becomes AgCl. This reaction releases an electron to the external circuit and leaves a free Flx^+^ cation. The Flx^+^ then migrates to the wound under electrophoretic force. The cathode reaction consumes an electron and releases a free Cl^-^, which can balance with the incoming cation. (C) A CAD model of the ion pump, made from PDMS. The ion pump has four reservoirs for fluoxetine solution, where electrode wires made from Ag and AgCl are inserted and connected to the external circuit through contact pins made of steel. The scale bar is 5 mm. (D) CAD models of the wearable bioelectronic bandage device assembly, with the controller module on top and the ion pump on the bottom, are displayed. The scale bar is 5 mm. (E) A camera photo of the wearable bioelectronic bandage is presented, and the scale bar is 5 mm. (F) The circuit diagram of the controller module is provided. It includes a microcontroller with a built-in clock and analog-to-digital converter (ADC), a memory chip, a digital-to-analog converter (DAC), and resistors for sensing current. (G) A camera photo of the ex-vivo test setup for the wearable bioelectronic bandage while the device is in contact with the saline solution inside a PDMS well is shown, and the scale bar is 5 mm. (H) The plot of the device’s current response to a 2V (peak to peak) square wave applied with a potentiostat is presented, indicating high temporal resolution.

Continuous delivery of Flx occurs due to two opposite electrochemical reactions at the working and counter electrodes that maintain charge balance throughout the system (Figure 2B). In the reservoir containing the working electrode or anode (Figure 2B, top), the protonated Flx is in Flx^+^Cl^-^ form to maintain charge neutrality. With V_Flx_ = 1-3 V, an electron leaves the surface of the Ag, creating Ag^+^, which in turn reacts with the Cl^-^ of the Flx·HCl. AgCl forms, leaving the positively charged Flx^+^ free to follow the electric field through the ion-selective hydrogel into the wound bed. To close the circuit, the opposite reaction occurs at the counter electrode (Figure 2B, bottom), where an electron from the leads creates Cl^-^ that pairs with one of the physiological cations, such as Na^+^ or K^+^, that entered the reservoir from the wound bed, leaving an Ag surface.

The ion pump module comprises four reservoirs and four contact pins that attach to the Ag/AgCl electrodes (Figure 2C). Two reservoirs deliver Flx^+^ into the wound bed, while the remaining two accept positive physiological ions to maintain charge balance in the wound bed. Comsol simulation results show that this delivery strategy provides a relatively uniform distribution of Flx^+^ on the wound (**Figure S3**). The ion pump module communicates with the control module through contact pins. The pins also provide mechanical coupling between the two modules (Figure 2D). On top of the ion pump module (Figure 2D and E), the control module contains microcontrollers, analog-to-digital converters, and a battery that makes it possible to function bandage wirelessly for a certain period of time. The control electronics translate the delivery regimen stored in memory into V_Flx_ pulses (Figure 2F). To test the ion pump delivery module, we disconnected the control module from the contact pins and directly connected the contact pins and electrodes to a potentiostat. We then actuated the ion pump module inside a well plate containing saline solution (Figure 2G). During this setup, we measured the current in the circuit (I_Flx_) while V_Flx_ underwent a series of pulses ranging from 0-2 V with a duration of 200 ms each. As expected, since the delivery circuit comprises all linear elements, I_Flx_ closely followed V_Flx_. According to our description of the delivery mechanism (Figure 2A and Figure 2B), I_Flx_ should directly measure the number of Flx^+^ molecules delivered to the wound bed, with each electron flowing into the circuit corresponding to an individual Flx^+^ molecule. To verify this hypothesis, we measured the amount of Flx^+^ delivered after a specific number of pulses into the saline solution (refer to SI text, **Figure S4**, **Table S1**). The HPLC results showed that the delivery efficiency (η) was 20±4%, meaning that we delivered one Flx^+^ molecule for every five electrons going through the circuit. This efficiency is in line with other ion pumps, and Flx^+^ is likely to face competition from H^+^ [15]. Calculating η allows us to infer the Flx^+^ dose from measuring I_Flx_ during delivery.

We then proceeded to test the efficacy of the bioelectronic bandage in vivo on a mouse wound model (**Figure 3**). We created a 6 mm punch biopsy wound held open by a silicone splint ring to minimize wound contraction (Figure 3A). Wound contraction is a major wound healing process in mice but not humans. It confounds the histological analysis of the wound reponse to therapy, particularly re-epithelialization [12b, 13]. A gas-permeable, transparent adhesive dressing, Tegaderm^TM^, fixes the bioelectronic bandage to the mouse (Figure 3A, right) so that the mouse can move around its cage without interference (Figure 3B). This is possible because the lightweight bandage (2.5 g) represents less than 10% of the mouse weight. For this test, we programmed the bandage to deliver Flx^+^ for three days for six hours per day (Figure 3C). From our estimates, this program should deliver approximately 100 nMol of fluoxetine per wound per day - a Flx^+^ dose that has been shown to improve healing in a mouse wound model when topically applied to the wound bed [12b, 13]. A blinking LED on the bioelectronic bandage indicated that the program was running as desired (**Figure S5**, **Movie S1**).

**Figure 3.**
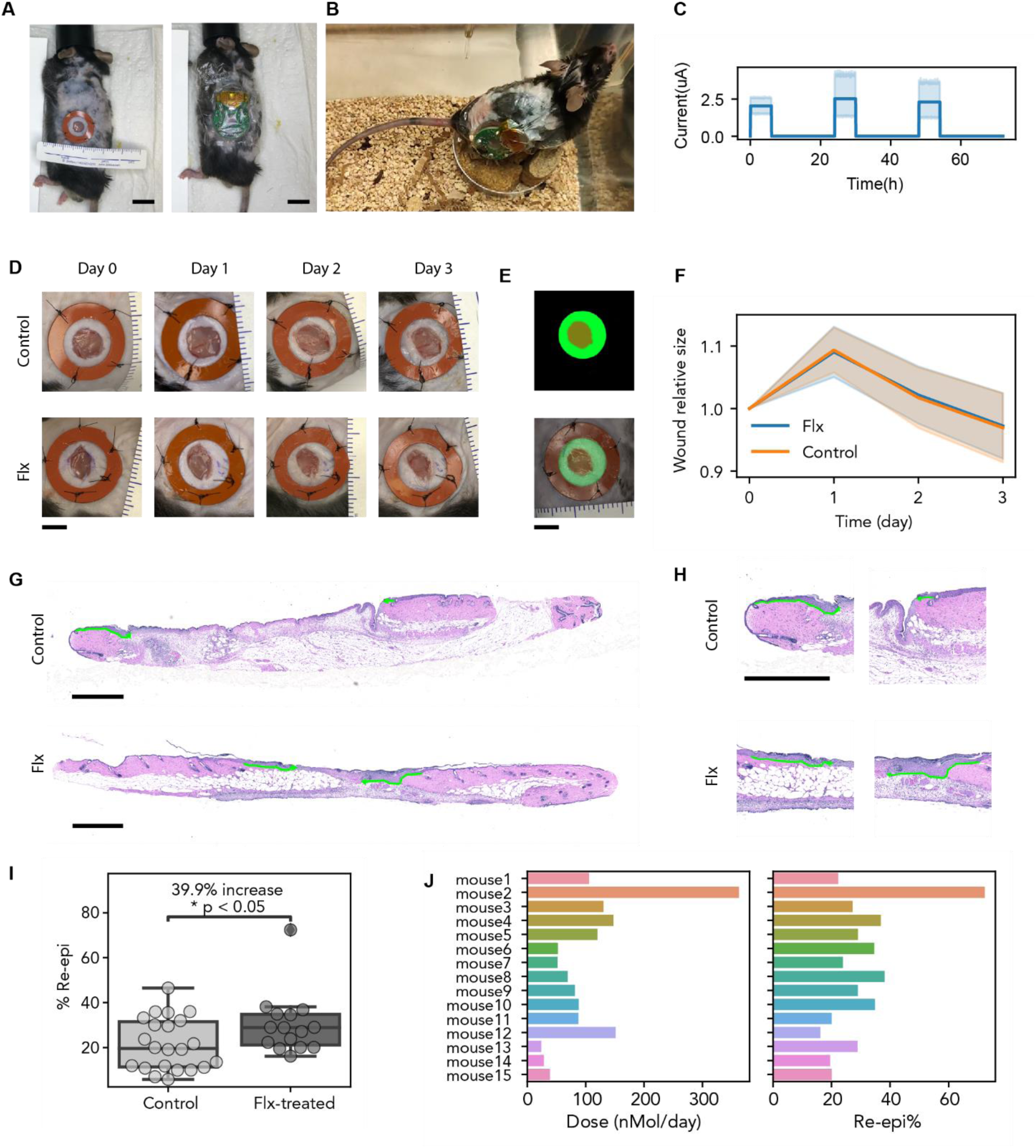
In vivo experiment of wearable bioelectronic bandage on mice. (A) Photo of a mouse wound model during surgery. The left panel shows the mouse wound sutured with a silicone splint ring, and the right panel shows the wearable bioelectronic bandage fixed on the wound by Tegaderm. Scalebar = 5 mm. (B) A mouse maintains normal daily activity while wearing the bioelectronic bandage on its wound. (C) Plot of fluoxetine actuation current over time from 15 experimental mice. (D) Wound image taken from a group of experiments over 3 days. Scalebar = 5 mm. (E) Image processing using machine learning to determine wound size. The top image is the mask generated by the algorithm to mark the size of the wound, and the bottom image depicts the overlayed image of the mask and the wound, showing that the masked area is accurate. (F) Wound area analysis of the control group (orange line) and fluoxetine-treated group (blue line). (G) H&E staining of tissue section across the wound. The top panel shows the control wound, and the bottom panel shows the fluoxetine-treated wound. The green line shows re-epithelization. Scalebar = 1 mm. (H) A zoomed-in view of the H&E staining on the area of the wound edge. Scalebar = 1 mm. (I) Box plot of statistics of the re-epithelization percentage of the wound. The control group has 21.6 +/-11.6% re-epithelization, and the fluoxetine-treated group has 30.2 +/-13.5% re-epi. Wounds treated with fluoxetine demonstrated a 39.9% increase in re-epithelization compared to the control, n=15-22 mice, p<0.05. (J) Bar plot of dose vs. re-epithelization percentage on each fluoxetine-treated mouse.

We utilized a machine-learning algorithm previously reported [16] to analyze images of wounds and evaluate their condition and healing progression (Figure 3D). The algorithm automatically marked the wound area and analyzed the wound size (Figure 3E). Both the control and fluoxetine-treated wounds exhibited a similar trend of a slight increase in size at day 1, followed by a decrease on subsequent days, which aligns with the expected size change during wound healing (Figure 3F). To further investigate healing, the wounds were fully excised at the end of the experiment, fixed, and stained with hematoxylin and eosin (H&E) to assess wound re-epithelialization (Figure 3G-H). Re-epithelialization, contributed to by both keratinocyte proliferation and migration, is required for wound closure [17]. Treatment of wounds with Flx^+^ delivered from the bioelectronic bandage resulted in a 39.9% (P<0.05) increase in re-epithelialization compared to control (Figure 3I), indicating a significant improvement in early-stage healing. Across 15 mice, the bioelectronic bandage consistently delivered the desired dose with some variability (Figure 3J, left). This variability in delivered dose can be easily amended with current control using closed-loop control algorithms, as we have previously demonstrated with other ion pumps [18]. This variability allowed us to qualitatively correlate the dose of Flx^+^ and wound healing as measured by the percentage of re-epithelization (Figure 3J, right). For example, re-epithelization in mouse 2, which received >300 nMol per day Flx^+^, was much faster than in mouse 15, which received a 10x smaller dose. This observation is consistent with the direct topical application of fluoxetine to murine skin wounds [12b, 13]. While the correlation between fluoxetine dose and re-epithelization is strong (R^2^ = 0.58), it is not statistically significant due to the small sample size (**Figure S6**).

After analyzing the promising re-epithelization data, we delved deeper into another crucial indicator of wound healing: the M1/M2 macrophage ratio (**Figure 4**) [19]. Macrophages play a pivotal role in the immune response and are crucially involved in the healing and regeneration of wounds. Although macrophages are phenotypically heterogeneous over a continuum, a simplified classification based on their polarized functions during the different stages of wound repair [19] defines the M1 subtype that carries out pro-inflammatory activities [20] and the M2 subtype that is anti-inflammatory [21] and promotes tissue repair [22]. Using immunohistochemistry to identify the subtypes based on the expression of recognized markers, we found M1 macrophages significantly infiltrated the center of the control wound three days after injury, while there was no change observed in the fluoxetine-treated wound (Figure 4A). Meanwhile, the number of M2-like macrophages increased in the center of the fluoxetine-treated wound, as depicted in Figure 4A. At day 3 the M1/M2 ratio was noticeably reduced following treatment, as shown in the overlay of M1 and M2 cells (Figure 4B). Fluoxetine treatment additionally reduced the M1/M2 ratio on day 3 by 27.2% compared to control (P<0.05) (Figure 4C) indicating a lower number of the M1 pro-inflammatory macrophages compared to the M2 pro-reparative macrophages. This M1/M2 ratio decrease suggests a shorter inflammatory phase with a more rapid progression towards the reparative phase of healing, consistent with the noted improvement in wound re-epithelization.

**Figure 4.**
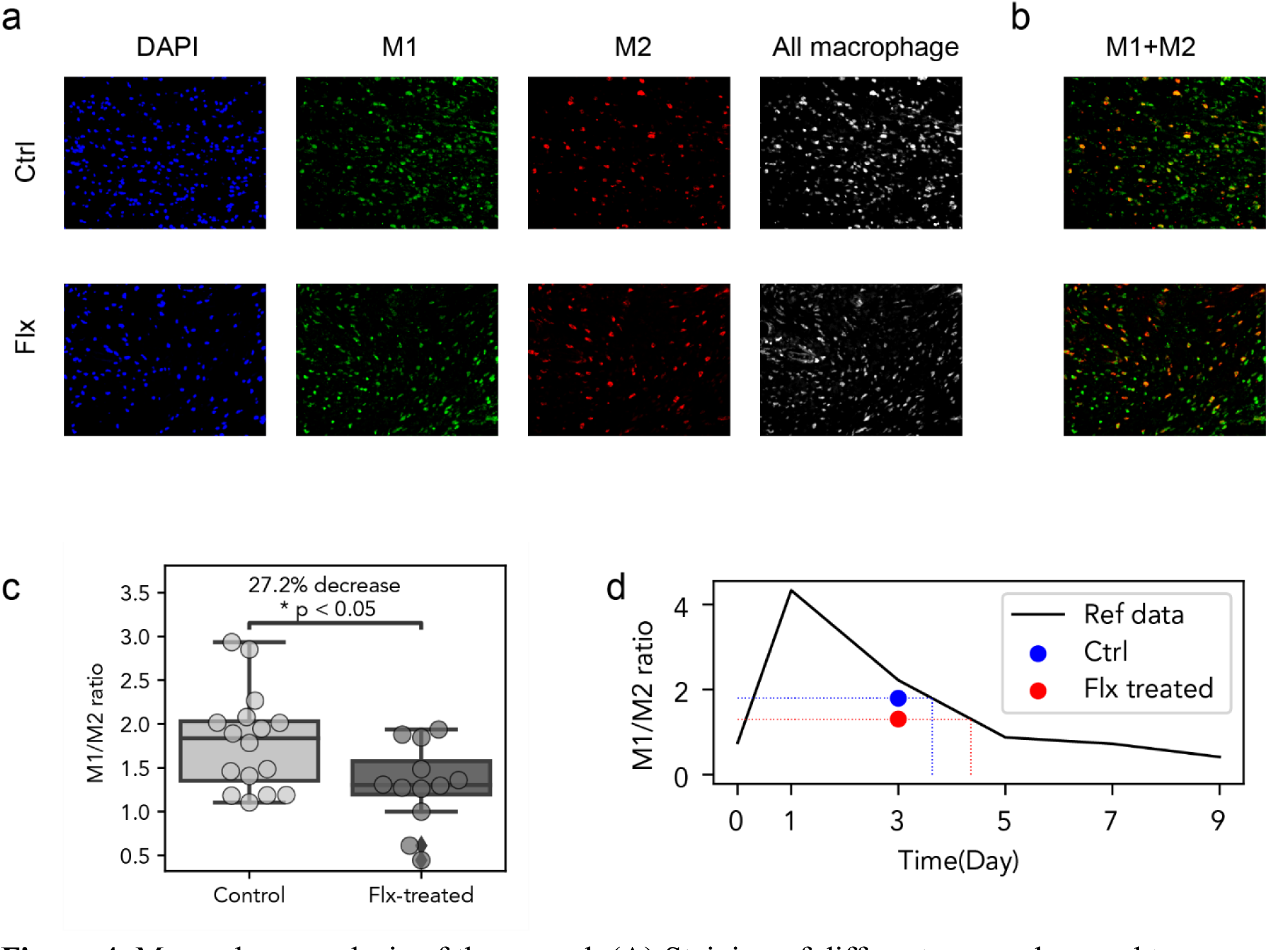
Macrophage analysis of the wound. (A) Staining of different macrophage subtypes in the wound section. All cells (stained by DAPI, shown in blue), M1 (stained by iNOS, shown in green), M2 (stained by CD206, shown in red), and overall macrophages (stained by F4/80, shown in gray) were imaged and presented. (B) Overlay of M1 and M2 staining, showing the distribution and ratio of macrophage subtypes. (C) Box plot showing statistics of the M1/M2 ratio, which was 1.80 +/-0.62 in the control group and 1.31 +/-0.46 for the fluoxetine-treated group. The fluoxetine-treated wounds demonstrated an average of 27.2% decrease in M1/M2 ratio compared to control wounds, n=13 mice, p<0.05. (D) Plot of the M1/M2 ratio over time. The blue dot represents the M1/M2 ratio from the control group, and the red dot represents the fluoxetine-treated group. The dashed lines indicate the projected wound age, while the solid line represents the trend plotted from the data in the reference. [23]

To further investigate this effect, we examined the M1/M2 ratio change in the context of a continuous curve over the healing process. Using time series data of M1 and M2 cells in mouse incision wounds obtained from published studies [23], we plotted the M1/M2 ratio’s dependence on time (Figure 4D). Though the comparison of the published data generated from incisional wounds [23] to the current data generated from excisional wounds may be an imperfect approach, nevertheless, plotting M1/M2 ratio along with published data enables us to estimate the wound progression. The M1/M2 ratio of both fluoxetine-treated and control wounds on day 3 of this experiment are within the same order of magnitude (blue and red dots in Figure 4D) as the time-series obtained from published data (black line in Figure 4D). However, because the curve is non-monotonic, there are two periods that might correspond to the M1/M2 value obtained in this study – one is day 0, and another is days 3-5. After taking into account other wound indicators, such as the onset of re-epithelialization, the presence of macrophages, and diminishing wound size, we conclude that our day 3 data correspond to the day 3-5 part of the curve, i.e. M1/M2 ratio is monotonically decreasing in that time period. Therefore, a lower M1/M2 ratio indicates that the wound has entered a later healing stage. Previous studies reported that the M1-M2 transition is critical for the resolution of inflammation and for promoting tissue repair [24]. Since the fluoxetine-treated wounds showed increased re-epithelialization and decreased M1/M2 ratio, we conclude that the wearable bioelectronic bandage’s fluoxetine treatment accelerated the wound-healing process. This finding is consistent with earlier studies that directly applied fluoxetine to the wound bed [12b, 13]. Although fluoxetine is a selective serotonin reuptake inhibitor (SSRI) primarily used as a systemically administered drug for treating depression, recent studies have revealed that SSRIs may affect various types of cells involved in cutaneous wound healing, such as keratinocytes, fibroblasts, endothelial, and immune cells [25], and modify their migration, differentiation, and function. Moreover, since the wearable bioelectronic bandage’s delivery is localized to the wound, there is little to no systemic accumulation of fluoxetine or impact on the serotonin metabolism (**Figure S7**), reducing unwanted side effects [26] (SI). As this research is a proof of concept for the in vivo application of wearable bioelectronic bandages for drug delivery, the sample size is small, and the treatment duration is short, causing the result to have wide distribution. Also, we only used the bioelectronic device to deliver a pre-determined dosage repetitively instead of delivering a customized treatment regimen. Though we proved that bioelectronic drug delivery helped wound healing compared to no treatment, it calls for more detailed studies to further compare it with other drug delivery methods such as topical application or conventional gradual release encapsulations to fully prove the advantage of the wearable bioelectronic bandages.

## 3 Discussion

Our results demonstrate the successful design and fabrication of a novel wearable bioelectronic bandage capable of actively controlling drug release for in vivo wound healing. The bandage is a standalone device that includes an ion pump delivery module, a controller module, and a power supply, eliminating the need for external instruments or connections to operate. This configuration allows for extended periods of use without external interaction. We created a wound treatment that provides the timed and programmed delivery of fluoxetine in a mouse wound model. Our treatment resulted in faster wound healing at day 3 as indicated by a statistically significant 39.9% increase in re-epithelization and 27.2% decrease in M1 pro-inflammatory macrophages respect to M2 pro-reparative macrophages. This suggests that fluoxetine treatment from the wearable bioelectronic bandage promotes wound healing by influencing the balance of M1/M2 phenotypes and cell regeneration. While the effect of fluoxetine accelerating wound healing is not surprising, the wearable bioelectronic bandage offers a customized drug dose regimen with temporal precision, which can be favorable over the conventional systemic routes of drug administration that deliver a single, full dose every 24 hours with an initial peak and subsequent lower concentration. In future research, if the timing of drug release can be programmed to align with the wound stage, it may further aid wound healing. Overall, the wearable bioelectronic bandage expands the application of bioelectronics on programmed drug delivery and demonstrates that the fluoxetine released from the wearable bioelectronic bandage retains its reparative biological activity. We aim to further optimize the design of the wearable bioelectronic bandage for potential clinical use in the future.

## 4 Materials and Methods

### 4.1 Experimental Design

The objective of this experimental research is to evaluate the effectiveness of a wearable bioelectronic bandage with an integrated electrophoretic ion pump for programmable drug delivery and enhanced wound healing in mice. We hypothesize that the bandage can deliver drugs with precise dose control and personalized therapeutic profiles to wounds in mice, leading to faster overall healing with minimal side effects.

The wearable bioelectronic bandage was tested in vivo using a mouse wound model. A total of 28 male C57B6 mice were used, with 15 in the treatment group and 13 in the control group. Six-millimeter wounds were created on the back of each mouse under anesthesia, using the protocol we have previously reported [12b]. The control group received a wearable bioelectronic bandage that was not powered on, while the treatment group received six hours of fluoxetine delivery per day for three days. The wound area, re-epithelialization rate, and M1/M2 macrophage ratio were evaluated using digital photography and histological analysis.

### 4.2 Design of wearable bioelectronic bandage

The wearable bioelectronic bandage was made up of two major components: the delivery module, which included an ion pump, and the controller module, which comprised a PCB, electronic components, and a battery. These components were fabricated separately and then assembled together.

### 4.3 Fabrication of ion pump

To fabricate the ion pump for the wearable bioelectronic bandage, AutoCAD software was used to design 3D-printed, two-part molds (Figure S1a). The molds were filled with Polydimethylsiloxane (PDMS) and baked at 60°C for 48 hours. The resulting PDMS parts were then removed from the molds (Figure S1b) and cleaned with Isopropyl Alcohol (IPA) and water, followed by nitrogen (N_2_) drying to ensure no debris remained on the PDMS layers. The top layer of PDMS contained four reservoirs, designed to hold fluoxetine solutions of specific concentrations, and four capillary tubes filled with hydrogels for fluoxetine delivery. The bottom PDMS part acted as a lid, covering the reservoirs and featuring a 0.5 mm tall notch to ensure contact with the wound bed below the skin. Silver (Ag) and silver-silver chloride (Ag/AgCl) wires with a diameter of 0.1mm were inserted inside each reservoir. The top and bottom layers were bonded together through oxygen (O_2_) plasma treatment, which oxidizes the polymer surface and changes the CH_3_ groups on the PDMS surface to OH groups. The oxidized surfaces were bonded together using custom aluminum pieces. The PDMS surface was coated with Parylene to increase the media lifetime on the PDMS reservoirs. Hydrogel-filled capillaries, which act as the ion exchange membrane for the ion pump, were fabricated using a previously optimized and reported method [27]. The hydrogel recipe in this study consisted of a 1 M concentration of 2-acrylamido-2-methyl-1-propanesulfonic acid (AMPSA), 0.4 M concentration of polyethylene glycol diacrylate (AMPSA), and 0.05 M concentration of photoinitiator (I2959). 100 mm of silica tubing with an inner diameter of 100 μm and an outer diameter of 375 μm were etched with NaOH and then treated with silane A174 to prevent hydrogel expansion. The hydrogel was crosslinked with five minutes of 365 nm UV treatment at a power density of 8 mW cm^-2^. After UV curing, the capillary tubes were segmented into 5 mm segments and loaded by immersing them in a 0.01 M fluoxetine solution for at least 4 hours before use. Finally, the capillaries filled with hydrogel were inserted into each reservoir to complete the fabrication of the ion pump.

### 4.4 Design and fabrication of controller module

The controller module for drug delivery comprises a programmable PCB with electronic components used for actuation and sensing. The PCB was designed using Autodesk EAGLE and contains a microcontroller with a built-in ADC, a memory chip, a DAC, and resistors for sensing current (Figure 2F). Prior to the experiment, programs were flashed into the onboard microcontroller, and actuation commences automatically once the battery is inserted. During program execution, the microcontroller sends I2C commands to the DAC to apply the appropriate voltages to the electrodes of the ion pump. As a result, current flows through the resistors, generating voltages that can be read by either the ADC on the microcontroller or by external probes.

### 4.5 Assembling of the wearable bioelectronic bandage

To integrate the two modules of the wearable bioelectronic bandage, steel pins were inserted into four holes on the PDMS layer of the ion pump. The bottom of each pin was coated with silver paste to establish electrical connections between the pin and Ag or Ag/AgCl electrodes. After assembling the wearable bioelectronic bandage, the pins were soldered to the PCB. Sterilized fluoxetine hydrochloride solutions (0.01 M) were then prepared by dissolving the drug in sterilized water, adjusting the pH to 6 to allow the fluoxetine to protonate, filtering through 0.2 µm filters, and injecting the sterilized fluoxetine solutions into each reservoir.

### 4.6 Ex vivo testing of the wearable bioelectronic bandage

The wearable bioelectronic bandage was tested ex vivo in PDMS wells filled with Steinberg solution to mimic the biochemical environment of tissue. The testing involved connecting the wearable bioelectronic bandage to a potentiostat (Metrohm Autolab) and controlling the voltage pattern via a computer while the ion pump outlet contacted the solution in the well. After a specific duration of actuation, voltage and current were recorded, and the test was stopped to collect the solution in the well. The total charge that went through the circuit was calculated by integrating the current over time, and the dose of fluoxetine was calculated by HPLC-MS. The efficiency of the delivery η= moles of fluoxetine/moles of charge.

### 4.7 Measurement of Fluoxetine with HPLC-MS

In order to measure the concentration of fluoxetine in the solution, we utilized HPLC-MS (Thermo Scientific™ LTQ) with a reversed-phase column (Synergi™ 4 µm Fusion-RP 80 Å, 150 x 2 mm, 00F-4424-B0). Initially, we generated a standard curve from a series of samples with known concentrations of fluoxetine. These samples were then analyzed using HPLC-MS to verify the fluoxetine peak by retention time and mass reading, and the mass-spec intensity was recorded. We plotted the peak area of mass-spec intensity vs. concentration in (Fig S4) and fitted the data to a calibration curve to obtain the slope and intercept. Subsequently, we loaded the collected samples with unknown concentrations into the HPLC-MS and recorded the fluoxetine intensity. Using the peak area and calibration curve, we were able to calculate the concentration of the sample.

### 4.8 Mouse preparation and drug delivery by the wearable bioelectronics bandage

We tested the bandage on C57B6 male mice by creating full-thickness, 6 mm circular wounds on their backs [12b]. We then applied wearable bioelectronics bandages loaded with either fluoxetine or a mock control device to the wounds and secured them with a Tegaderm overwrap. The target delivery dose for fluoxetine was set at 100 nMol or 0.035 mg/wound/day during the daily 6-hour delivery program. On the third day after the surgery, we removed the wearable bioelectronics bandages, captured images of the wounds using a camera, and harvested wound tissue for histological analysis.

### 4.9 Tissue collection and sectioning

After euthanizing the mice at the end of each experiment, wounds were excised and placed in a paraformaldehyde solution for fixation for 24 hours. Fixed tissues were then processed in a tissue processor for formalin-fixed paraffin-embedded (FFPE) tissue histology. During processing, the tissues were dehydrated and impregnated with paraffin wax to preserve the tissue structure. The processed tissues were embedded into paraffin blocks and cut into 5-µm thick sections using a microtome, and the sections were placed onto glass slides. After drying, these sections were used for further analysis of re-epithelialization and macrophage subtypes.

### 4.10 Re-Epithelization Calculation

To assess re-epithelialization, tissue sections were stained with H&E using a standard protocol. Brightfield images were taken on a BioRevo BZ-9000 inverted microscope, and BZ Analyzer software (Keyence, Osaka, Japan) was used to score the images. The left and right wound edges were determined by identifying the innermost follicle on each side. The basal keratinocyte layer was used to measure epithelial ingrowth on each side of the wound, from the innermost follicle to the tip of the epithelial tongue. The total width of the wound was measured along the surface of the granulation tissue between the two follicles. The percent re-epithelialization was calculated as [(length of left epithelial tongue in µm) + (length of right epithelial tongue in µm)]/(wound width in µm) X 100.

### 4.11 Macrophage Staining

We collected full-thickness mouse skin tissues from both fluoxetine-treated and untreated groups three days after wounding and examined them for the expression of M1 (F4/80+iNOS+) and M2 (F4/80+CD206+) macrophage markers. To detect all macrophages, we used F4/80 as a pan-macrophage marker. We identified M1-like macrophages as a fraction of F4/80+ macrophages expressing inducible nitric oxide synthase (iNOS), and M2-like macrophages as F4/80+ cells expressing CD206.

Formalin-fixed, paraffin-embedded tissues were deparaffinized, processed for antigen retrieval, and blocked for 2 hours at room temperature with 10% Donkey Serum (Thermo Fisher). The slides were then incubated overnight at 4 °C with primary antibodies, including Rat anti-F4/80 (dilution 1:50; MCA497G, BIO-RAD, Hercules, CA), Rabbit anti-iNOS (dilution 1:100; PA3-030A, Thermo Fisher Scientific), and Goat anti-CD206 (dilution 1:100; PA5-46994, Thermo Fisher Scientific). We selected these antibodies based on the validation information provided on the manufacturers’ websites.

After washing, sections were incubated with corresponding Alexa Fluor-conjugated secondary antibodies (Donkey Anti rat-AlexaFluor 488, Donkey Anti rabbit-AlexaFluor 647, Donkey Anti goat-AlexaFluor 568, dilution 1:200, Thermo Fisher Scientific). Nuclei were counterstained with 4’,6-diamidino-2-phenylindole (DAPI), and coverslips were mounted using an anti-fade mountant (SlowFade Mountant; S36936, Thermo Fisher Scientific). Fluorescence images were acquired using a Keyence automated high-resolution microscope (BZ-X800, KEYENCE, Itasca, IL). We imaged 5 adjacent areas at the wound center at 40x magnification and processed the images using ImageJ and CellProfiler 4.2 software.

For semi-quantification, we manually counted the numbers of macrophage subtypes in the wound based on double-positive staining with F4/80+iNOS+ for M1-like macrophages and F4/80+CD206+ for M2-like cells, with blind evaluation.

### 4.12 Fluoxetine quantitative analysis in blood

To measure the levels of fluoxetine and its major metabolite norfluoxetine in mouse serum, we employed reverse-phase HPLC with UV detection and purified the samples using C18 pipette-tip solid phase extraction (C18 PT-SPE). The mobile phase used was 300:700:1 acetonitrile: 50 mM phosphate buffer pH = 6.0: triethylamine. Separation of fluoxetine, norfluoxetine, and fluvoxamine (the internal standard) was achieved on a Cortecs C18 column (3 mm ID x 100 mm L, 2.7 um particle diameter, Waters, Ireland) at a flow rate of 450 uL/minute and a temperature of 35°C. The column effluent was monitored at 230 nm with a time constant of 0.5 seconds. For the PT-SPE cartridges, 20 mg C18 silica sorbent was slurry-packed in methanol into 1 mL pipette tips using glass wool as a bottom frit, and acid-washed sand in place of a top frit. After packing, we passed 2 tube volumes of water through and the cartridges were ready for use. To prepare the sample loading solution, we spiked 10 uL of a fluvoxamine solution (10 µg/mL in methanol) into 100 µL of study serum, and then diluted this solution with 100 µL of water. This sample loading solution was applied to the conditioned SPE cartridges, which were then washed with 600 µL ultra-pure water followed by 600 µL 50:50 methanol:water. We eluted fluoxetine, norfluoxetine, and fluvoxamine using 100 µL of methanol containing 0.5% (v/v) formic acid. After elution, we diluted the samples 1:1 with water, mixed well, and centrifuged them. Finally, we injected 10 µL of the resulting solution for analysis.

### 4.13 Serotonin quantitation in blood

Serotonin quantitation in mouse serum was performed according to our previously published protocol [28].

### 4.14 Statistical Analysis

Statistical analysis was performed using multiple software packages including Microsoft Excel, the Scipy.stats library written in Python, and Prism 9. The significance of the comparison of re-epithelialization and M1/M2 ratio between the treated and control group was assessed using an independent two-tailed Student’s t-test, with a significance level of P < 0.05. The correlation between the dose and re-epithelialization is evaluated by Pearson correlation coefficient (r), where r^2^ > 0.5 is considered a strong correlation. The significance of the linear regression was evaluated by Wald Test with t-distribution, with a significance level of P < 0.05.

## Supporting Information

Supporting Information is available from the Wiley Online Library or from the author.

## Acknowledgements

We thank the help from Mr. Vincent Pham and Dr. Harrison Shawa for performing the mouse experiments and animal care.

## Author contributions

Conceptualization: HL, HY, MT, RI, MR, JS

Methodology: HL, HY, PB, HH, TN, CH, KZ, JS, LL, HC, MT, MA, LL

Investigation: HL, HY, NA, PB, AL, AG, KZ, CR Visualization: HL, KZ, HC

Supervision: RI, MR, MT, MZ, MG, ML, AS, NN

Writing—original draft: HL, HY

Writing—review & editing: HL, HY, NA, TN, KZ, MT, MZ, MG, RI, MR

## Data and materials availability

All data are available in the main text or the supplementary materials.

