## supplemental file for "Programmable delivery of fluoxetine via wearable bioelectronics for wound healing in vivo"

**This PDF file includes:**

Supplementary Text
Figures. S1 to S7
Tables S1 to S2

**Other Supplementary Materials for this manuscript include the following:**
Movie S1

**Supplementary Text**

Measurement of ion pump efficiency

To calculate the efficiency of the ion pump, we divided the number of fluoxetine molecules transported across the membrane by the amount of charge passing through the circuit. We measured an average charge of 1.3e-3 C in the circuit after 10 minutes of ion pump operation. Next, we collected the saline solution from the PDMS well and used HPLC to measure the concentration of fluoxetine. Our analysis of five collected samples showed an average fluoxetine concentration of 138 uM in a 100 uL solution in the well. This corresponded to 13.8 nanomoles of fluoxetine and suggested an estimated 20% efficiency in converting charge to ion transport.

Measurement of systemic fluoxetine and serotonin Levels

To assess the overall Impact of the treatment, we measured the serum levels of fluoxetine and its metabolite norfluoxetine. We found that both were well below the physiological active level, with fluoxetine levels < 20 ng/mL and norfluoxetine levels < 50 ng/mL (except for one animal at 66.0 ng/mL). Furthermore, there was no significant difference in the level of serotonin in the blood between the control and Flx-treated animals (27.0 uM in control vs 23.0 uM in Flx-treated animals, p=0.36, detection limit 0.35 uM). These results suggest that the extremely low concentration of serum fluoxetine observed does not cause any systemic effects (52-54). HPLC Serotonin measurements show that there is no significant difference on serotonin concentration in the serum between the control and fluoxetine-treated group (Fig S7).

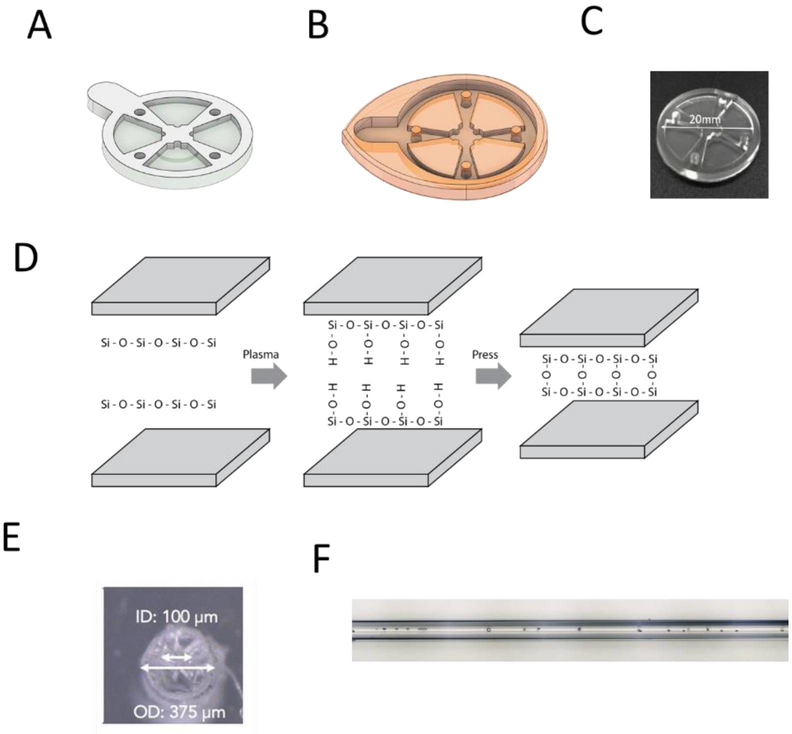


**Figure S1.**

Fabrication of the ion pump. (A) A CAD model of the ion pump body made from PDMS. (B) A CAD model of the casting mold for the ion pump body. (C) An optical picture of the casted PDMS piece. (D) A chemical process illustrating the plasma-assisted bonding of two PDMS pieces to fabricate the ion pump. The plasma activates the Si-Ox surface and exposes the hydroxyl groups, which bond with each other after pressure is applied. (E) A microscopy image of the cross-section of a capillary filled with AMPSA:PEGDA hydrogel. (F) A side view of the cross-section of a capillary filled with AMPSA:PEGDA hydrogel.

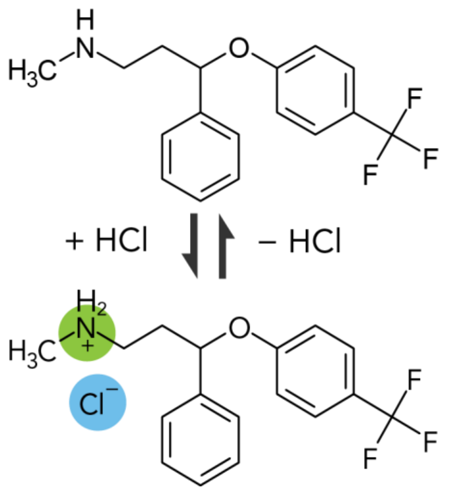


**Figure S2.**

Reversible protonation of fluoxetine with the presence of HCl. Fluoxetine combines with the $H^+$ and become $Flx^+$ cation at low pH.

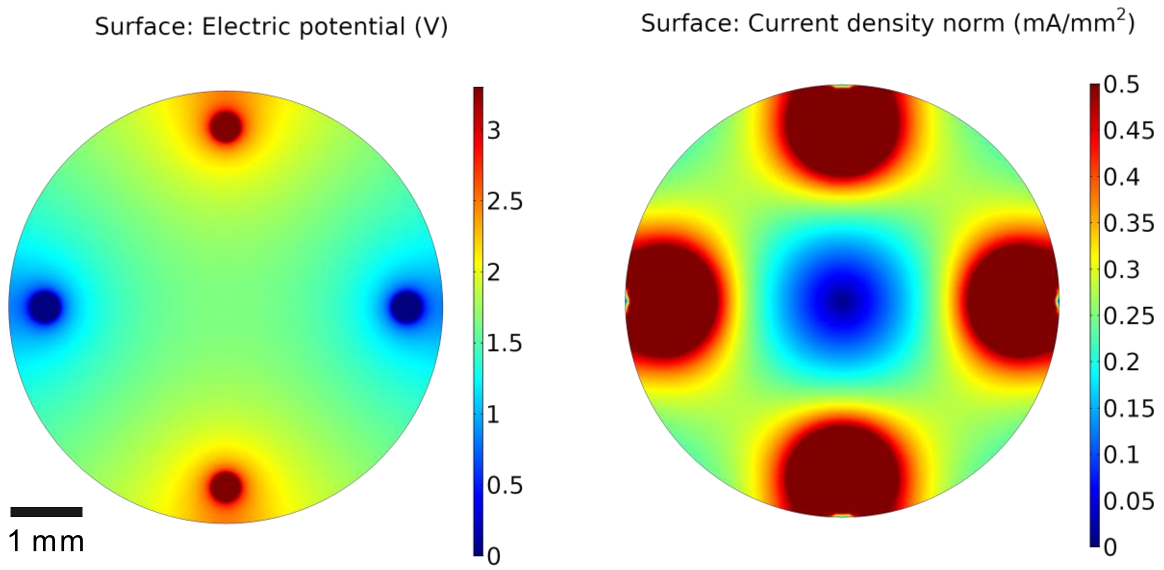


**Figure S3.**

Comsol simulation showing the distribution of Electrical Potential(left) and Current Density(right) when the wearable bioelectronic bandage operates.

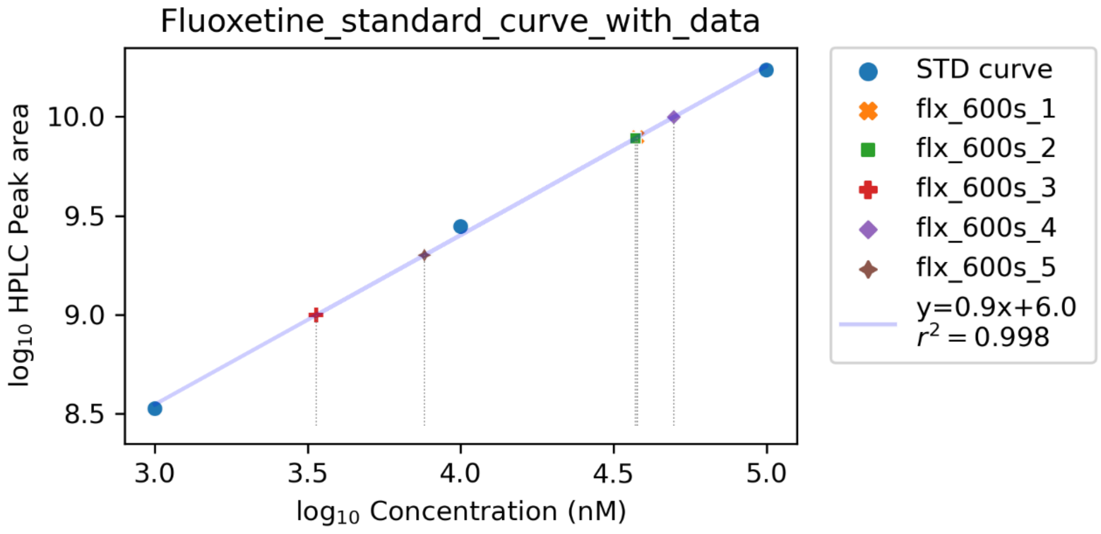


**Figure S4.**
HPLC-MS measurement of fluoxetine deliver from the wearable bioelectronic bandage tested ex vivo

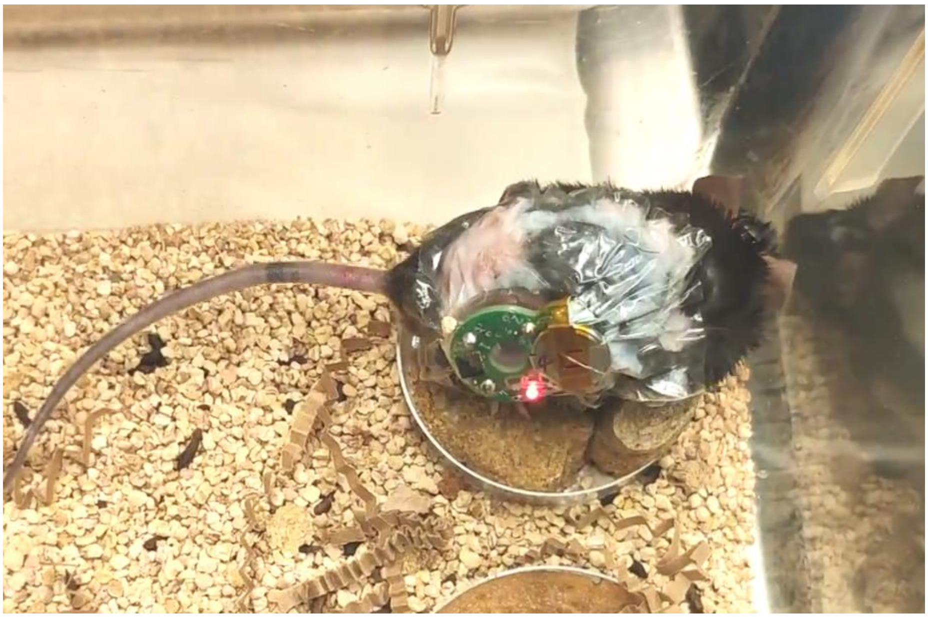

**Figure S5.**
LED blinking while the mouse live in the cage with the wearable bioelectronic bandage

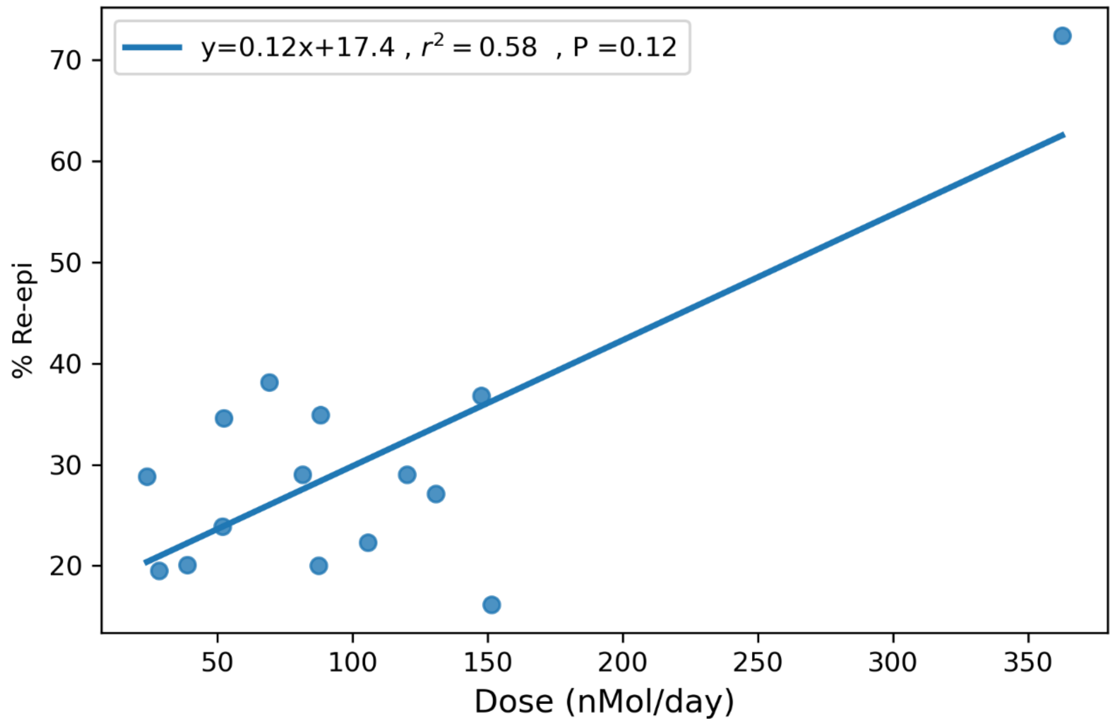


**Figure S6.**
Linear fitting of daily dose to re-epithelialization percentage. Square of Pearson correlation coefficient is 0.58. P value from Wald Test is 0.12.

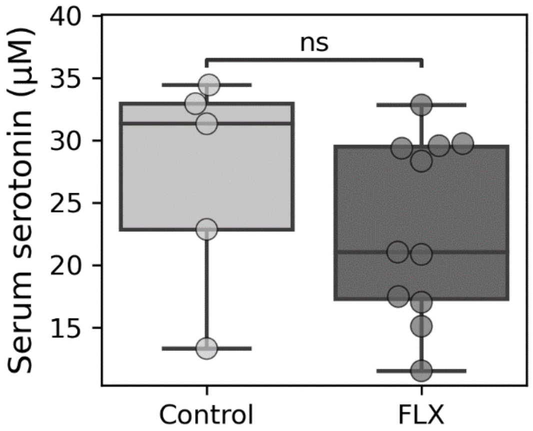


**Figure S7.**
Serum serotonin concentration of control and fluoxetine treated mice. Systemic serotonin concentrations did not differ significantly between the fluoxetine treated and untreated mice.

| Sample | Flx Conc(nMol/L) | Solution (μL) | Moles of Flx (nMol) | Charge (C) | Moles of electron (nMol) | Efficiency |
|---|---|---|---|---|---|---|
| Flx_600s_1 | 3.79E+04 | 150 | 5.68 | 2.76E-03 | 2.86E+01 | 19.9% |
| Flx_600s_2 | 3.73E+04 | 150 | 5.59 | 2.10E-03 | 2.18E+01 | 25.7% |
| Flx_600s_3 | 3.37E+03 | 300 | 1.01 | 6.37E-04 | 6.60E+00 | 15.3% |
| Flx_600s_4 | 4.98E+04 | 150 | 7.47 | 3.12E-03 | 3.23E+01 | 23.1% |
| Flx_600s_5 | 7.61E+03 | 150 | 1.14 | 6.60E-04 | 6.84E+00 | 16.7% |

**Table S1.**

Summary of ion pump efficiency calculated by charge in circuit and concentration measured from HPLC.

| Mouse | Mean current (μA) | Total charge per day (C) | Drug dose per day (nMol) | Drug dose per day (mg) | Re-epi (%) |
|---|---|---|---|---|---|
| 1 | 2.36 | 5.09E-02 | 105.7 | 0.033 | 22.3 |
| 2 | 8.08 | 1.75E-01 | 362.5 | 0.112 | 72.4 |
| 3 | 2.91 | 6.29E-02 | 130.7 | 0.040 | 27.1 |
| 4 | 3.29 | 7.11E-02 | 147.6 | 0.046 | 36.8 |
| 5 | 2.68 | 5.79E-02 | 120.1 | 0.037 | 29.0 |
| 6 | 1.17 | 2.52E-02 | 52.3 | 0.016 | 34.6 |
| 7 | 1.16 | 2.50E-02 | 52.0 | 0.016 | 23.9 |
| 8 | 1.54 | 3.33E-02 | 69.2 | 0.021 | 38.1 |
| 9 | 1.82 | 3.93E-02 | 81.6 | 0.025 | 29.0 |
| 10 | 1.96 | 4.24E-02 | 88.1 | 0.027 | 34.9 |
| 11 | 1.95 | 4.21E-02 | 87.4 | 0.027 | 20.0 |
| 12 | 3.38 | 7.29E-02 | 151.4 | 0.047 | 16.2 |
| 13 | 0.53 | 1.15E-02 | 23.9 | 0.007 | 28.9 |
| 14 | 0.63 | 1.36E-02 | 28.3 | 0.009 | 19.5 |
| 15 | 0.86 | 1.86E-02 | 38.7 | 0.012 | 20.1 |

**Table S2.**

Summary of dose and re-epithelialization percentage on 15 fluoxetine-treated mice.

**Movie S1.**

Provided in a separate file, showing the mouse moving around while wearing the wearable bioelectronic bandage on the back. LED is blinking, indicating the device is actively delivering fluoxetine.
